## Supplemental Figures for "Transcriptome-wide Association Study and eQTL colocalization identify potentially causal genes responsible for bone mineral density GWAS associations"

**Supplemental Figure 1. LSBMD and FNBMD GWAS SNPs in the PPP6R3 locus. A**) GWAS SNPs for LSBMD in the *PPP6R3* locus. SNPs in red are significant *PPP6R3* eQTL in thyroid tissue. The dashed line represents the genome-wide significance level (P-value= 5 x10$^{-8}$). **B**) GWAS SNPs for FNBMD in the *PPP6R3* locus. SNPs in red are significant *PPP6R3* eQTLin thyroid tissue. The dashed line represents the genome-wide significance level (P-value= 5 x10$^{-8}$). **C**) Mirroplot of LSBMD SNPs and *PPP6R3* eQTL. SNPs are colored by their LD with rs10047483 (purple), the most significant *PPP6R3* eQTL in thyroid. In the LSBMD panel, the most significant SNP is highlighted in purple , as rs10047483 was not assayed. **D**) Mirroplot of FNBMD SNPs and *PPP6R3* eQTL. SNPs are colored by their LD with rs10047483 (purple), the most significant *PPP6R3* eQTL in thyroid. In the FNBMD panel, the most significant SNP is highlighted in purple , as rs10047483 was not assayed.

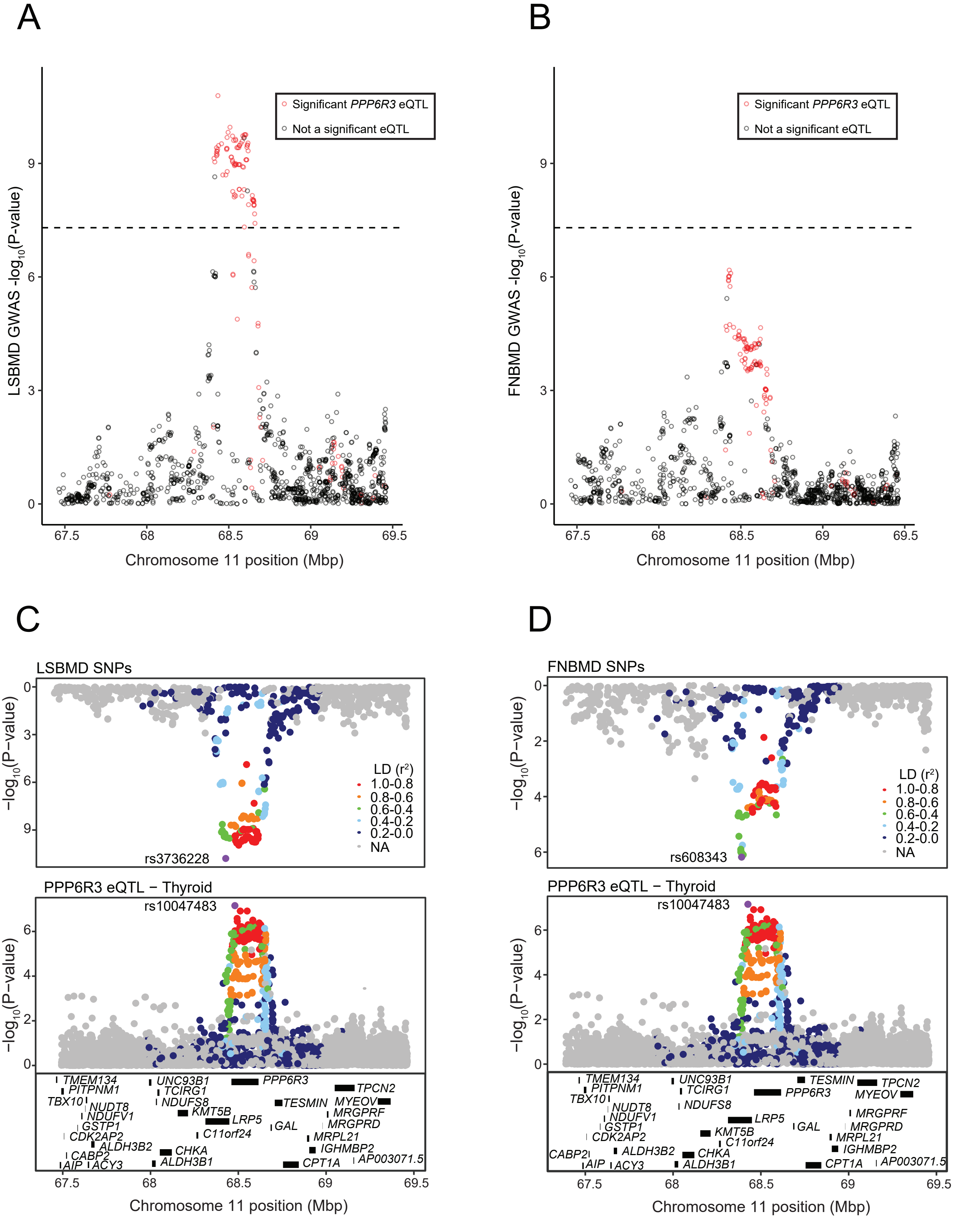

**Supplemental Figure 2. *Ppp6r3* functional validation. A**) Weight. B) Anterior-posterior (AP) femoral width. C) Medial-lateral (ML) femoral width. D) Femoral length (FL). E) Tissue mineral density (TMD), as measured by µCT. In all panels, least-square means are plotted. P-values are contrast P-values, adjusted for multiple comparisons. Asterisks represent significance (P<=0.05).

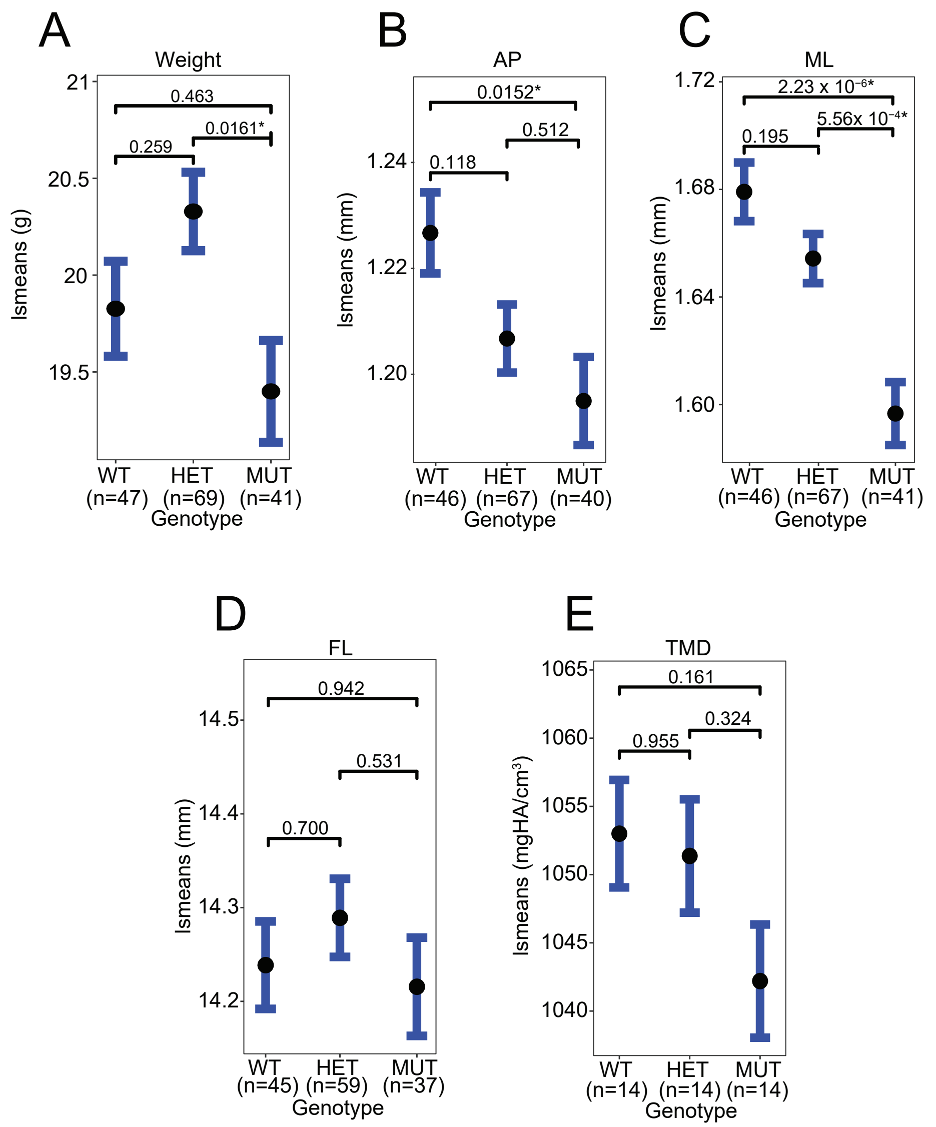

**Supplemental Figure 3. Raman spectroscopy in femur, means.** A-C) Least-square means for mean(mineral:matrix) in sex-combined, female, and male samples, respectively. D-F) Least-square means for mean(carbonate:phosphate) in sex-combined, female, and male samples, respectively. G-I) Least-square means for mean(crystallinity) in sex-combined, female, and male samples, respectively. Contrast P-values, adjusted for multiple comparisons, are presented.

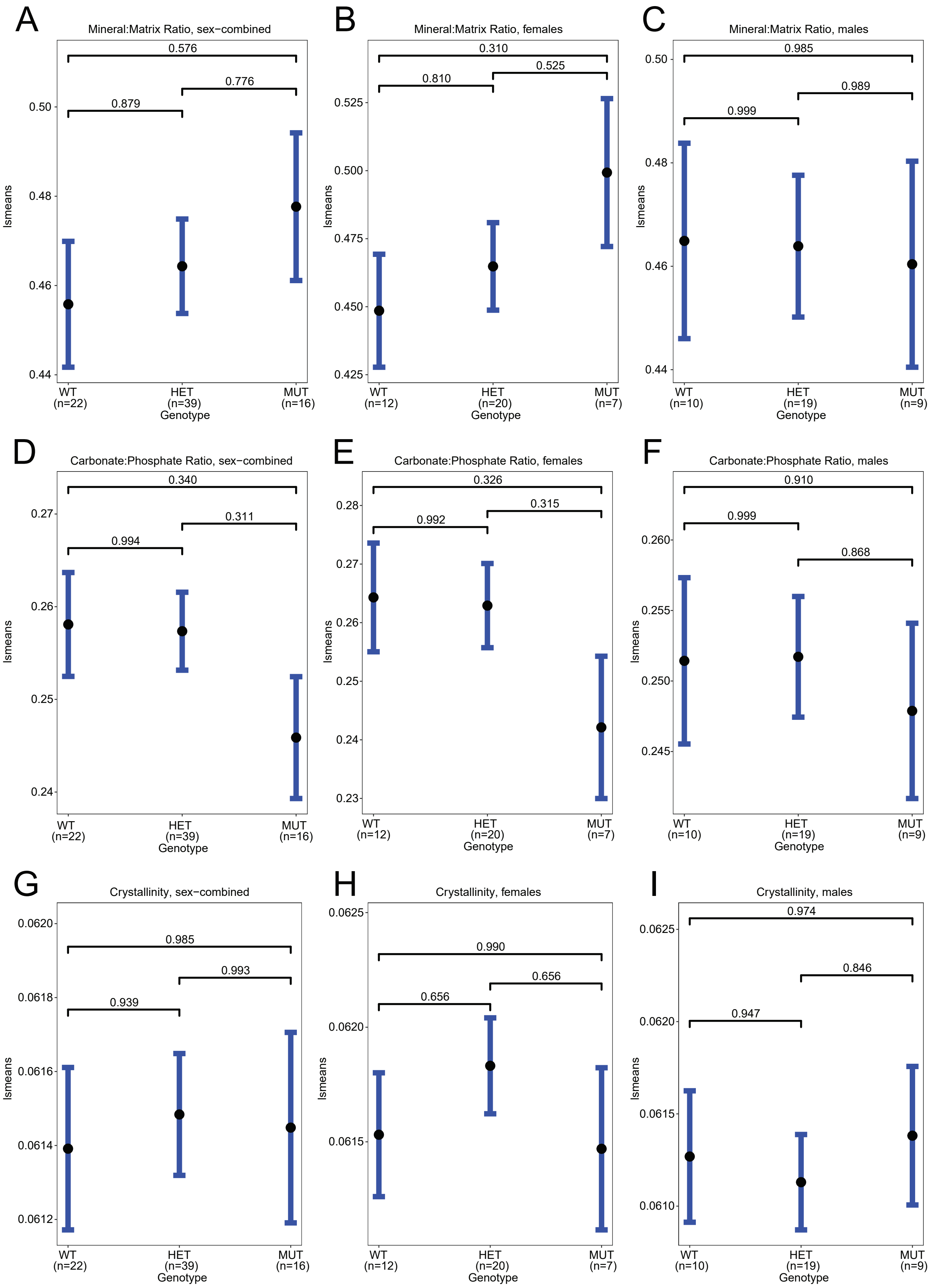

**Supplemental Figure 4. Raman spectroscopy in spines, means.** A-C) Least-square means for mean(mineral:matrix) in sex-combined, female, and male samples, respectively. D-F) Least-square means for mean(carbonate:phosphate) in sex-combined, female, and male samples, respectively. G-I) Least-square means for mean(crystallinity) in sex-combined, female, and male samples, respectively. Contrast P-values, adjusted for multiple comparisons, are presented.

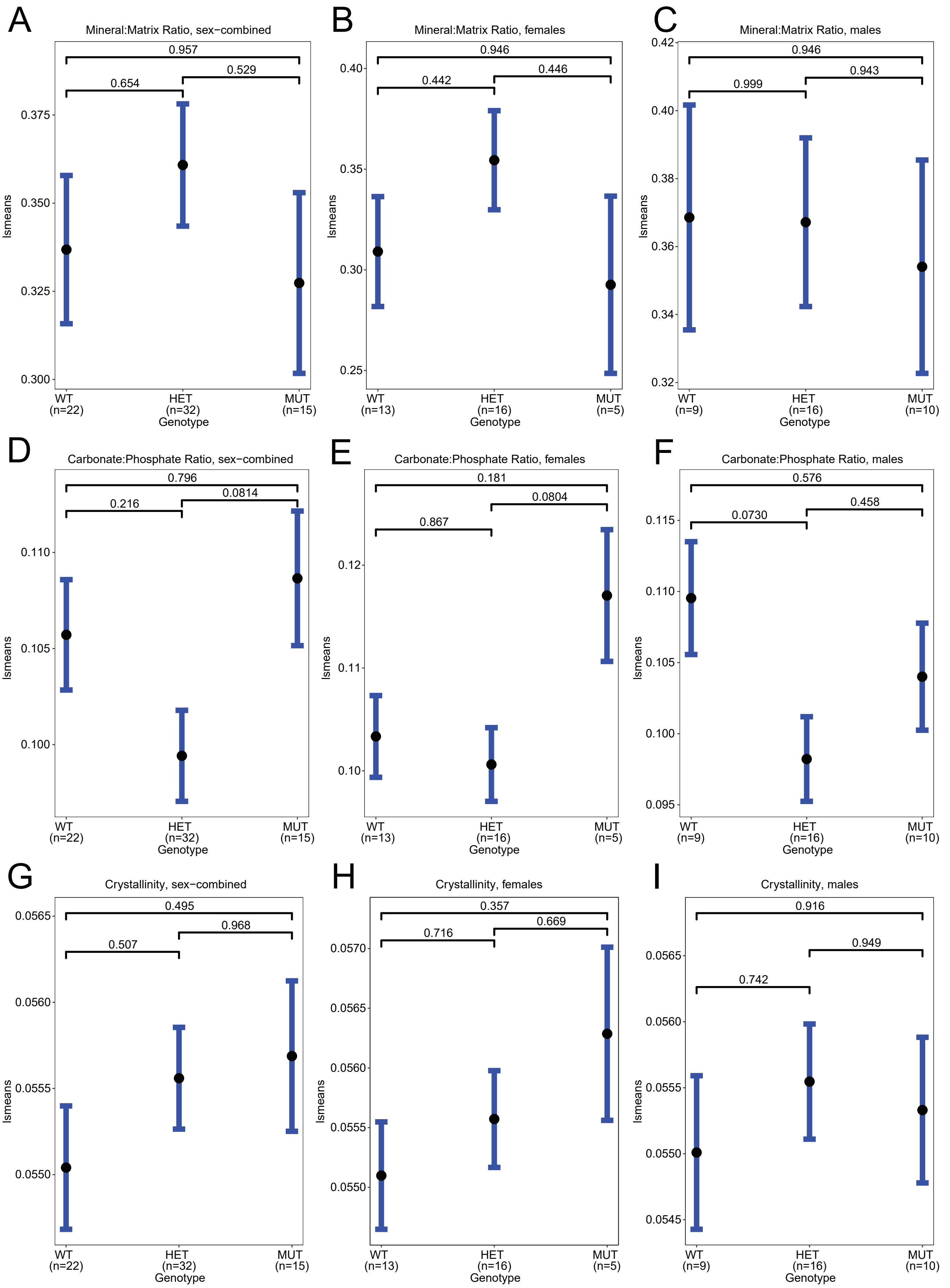

**Supplemental Figure 5. Raman spectroscopy in femurs, standard deviation.** A-C) Least-square means for sd(mineral:matrix) in sex-combined, female, and male samples, respectively. D-F) Least-square means for sd(carbonate:phosphate) in sex-combined, female, and male samples, respectively. G-I) Least-square means for sd(crystallinity) in sex-combined, female, and male samples, respectively. Contrast P-values, adjusted for multiple comparisons, are presented.

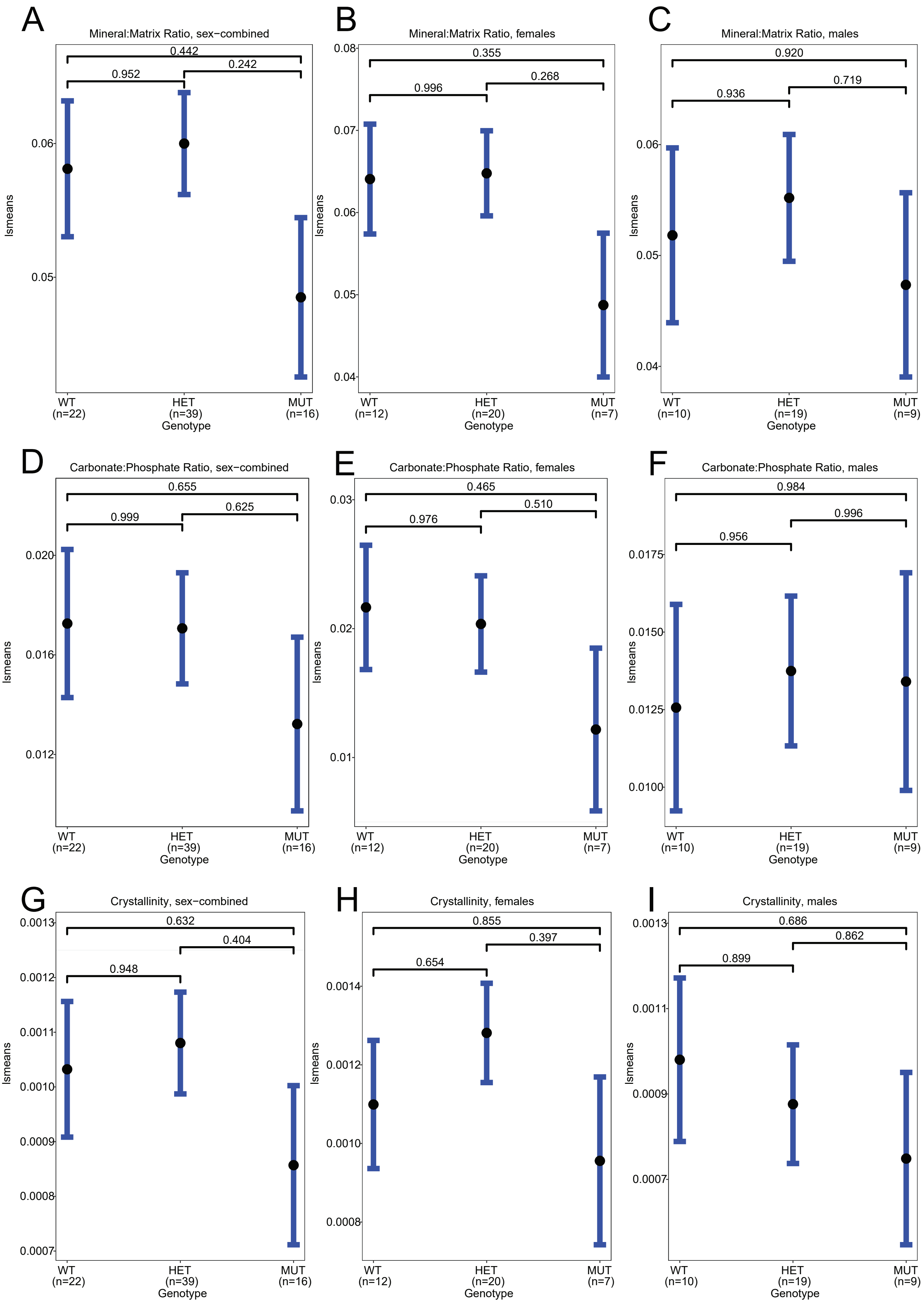

**Supplemental Figure 6. Raman spectroscopy in spines, standard deviation.** A-C) Least-square means for sd(mineral:matrix) in sex-combined, female, and male samples, respectively. D-F) Least-square means for sd(carbonate:phosphate) in sex-combined, female, and male samples, respectively. G-I) Least-square means for sd(crystallinity) in sex-combined, female, and male samples, respectively. Contrast P-values, adjusted for multiple comparisons, are presented.

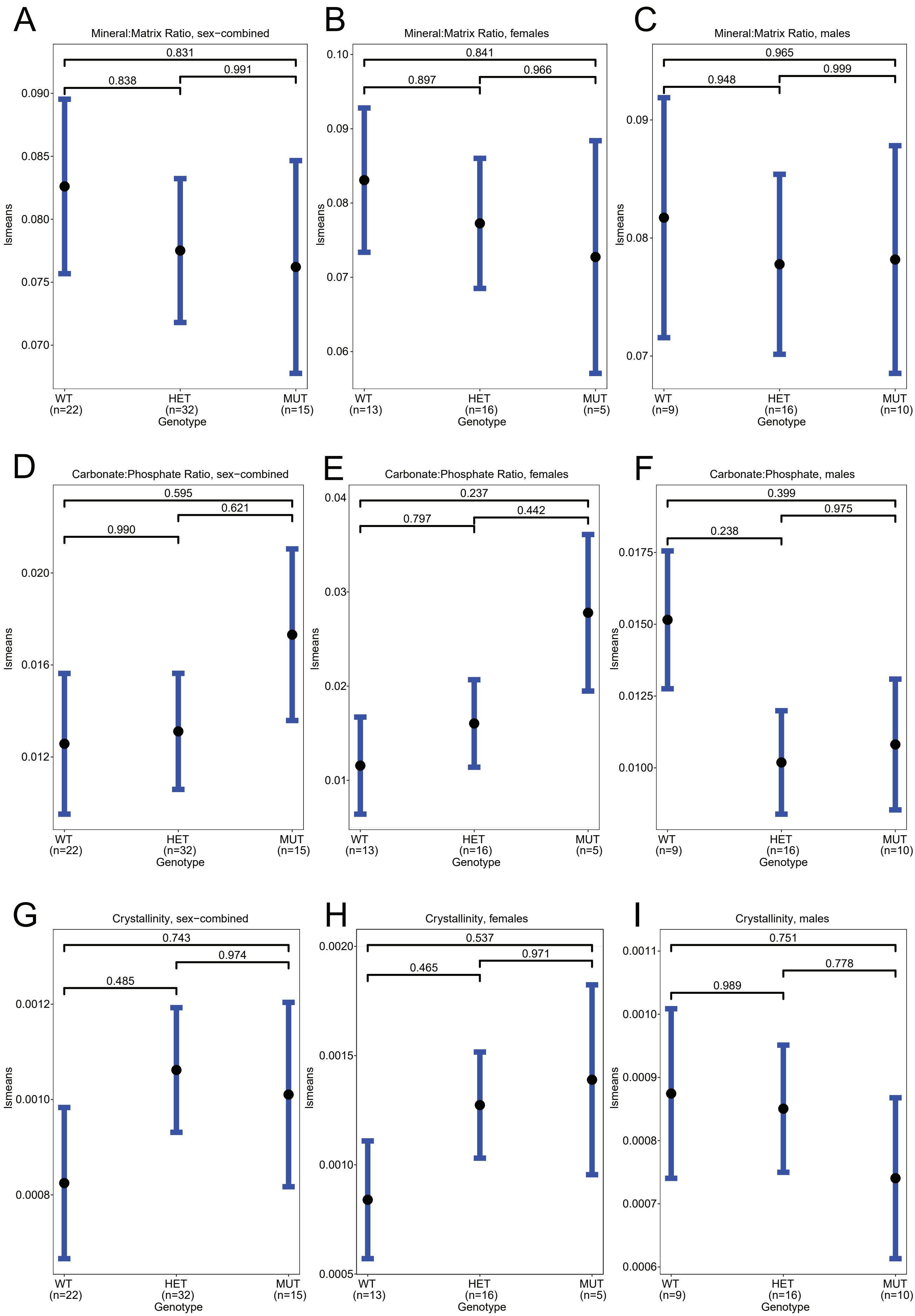
